# Onlinemeta: A Web Server for Meta-Analysis Based on Shiny Framework

**DOI:** 10.1101/2022.04.13.488126

**Authors:** Weiyuntian Dai, Yonglin Yi, Anqi Lin, Chaozheng Zhou, Jian Zhang, Shixiang Wang, Min Sun, Peng Luo

**Author notes:** Correspondence: **Shixiang Wang**; Address: Department of Biomedical Informatics, School of Life Sciences, Central South University, Changsha, PR China., **Min Sun**; Address: Department of General Surgery, Taihe Hospital, Hubei University of Medicine, Shiyan, China., **Peng Luo**; Address: Department of Oncology, Zhujiang Hospital, Southern Medical University, Guangzhou, 510282, Guangdong, China. Joint Authors. These authors have contributed equally to this work and share first authorship.

## Abstract

Meta-analysis is a common statistical method used to summarize multiple studies that cover the same topic. It can provide less biased results and explain heterogeneity between studies. Although there exists a variety of meta-analysis softwares, they are rarely both convenient to use and capable of comprehensive analytical functions. As a result, we are motivated to establish a meta-analysis web tool called Meta-Analysis Online (Onlinemeta). Onlinemeta includes three major modules: risk bias analysis, meta-analysis and network meta-analysis. The risk bias analysis module can produce heatmaps and histograms, whereas the meta-analysis module accepts many types of data as input, including dichotomous variables, single-armed dichotomous variables, continuous variables, single-armed continuous variables, survival data, deft method and diagnostic experiments, and outputs well-tuned forest plots, sensitivity analysis forest plots, funnel plots, comparison table of effects, SROC curve, and crosshair plots. For network meta-analysis module, the tool can process dichotomous variables or continuous variables, and generate network plots, forest plots, SUCRA (Surface Under the Cumulative Ranking) plots, rank plots and heatmaps. Onlinemeta is available at https://smuonco.Shinyapps.io/Onlinemeta/.

## Introduction

Meta-analysis is a statistical analysis strategy that synthesizes the results of independent studies covering the same topic to estimate the overall effect of the research, the stability of the results, and identifies any influencing factors[1, 2] . When evidence from multiple single studies is combined, a broad meta-analysis can solve problems such as compensating for insufficient sample size, reducing the risk of single-study bias, and providing more objective and effective results. Thus, the results of this type of analysis may help explain heterogeneity among studies and provide a direction for future research[3-5]. As new researches and data collection continue to accumulate, meta-analysis has provided researchers with an effective method to summarize these results and generate evidence-based medical data, has been widely used as a result. A search with filter ‘(“meta-analysis” or “meta-analysis”)’ in PubMed shows that from 2010 to 2024, the number of articles published using meta-analysis techniques is increased from 6993 to 35506 per year.

However, to our knowledge, many existing meta-analysis software have rooms for improvement. At present, the most commonly used meta-analysis software includes R, Stata, RevMan, and MetaXL. RevMan and MetaXL provide user-friendly interfaces, but they have limits in their statistical functions and graphic customization capabilities. In addition, the initial software setup is complicated, and the software must be downloaded separately[6, 7]. Stata is one alternative, but it is a commercial software which can be inconvenient to use. In addition, Stata analysis struggles when study outcome data has missing value and small-study effects appear[8]. R is free to use and has excellent statistical and graphic customization functions, but it requires users to be familiar with programming, thus limits the user population[9]. Therefore, we are motivated to develop a new tool with the following characteristics: (1) a user-friendly interface, requiring no specific programming knowledge; (2) no need to download and config; (3) non-commercial and free to use; (4) featuring with a one-stop functionality, capable of multiple meta-analysis types including network meta-analysis, meta-analysis for single-arm variable, deft method and diagnostic tests; (5) supporting highly customizable graphics.

With this in mind, we have developed the online meta-analysis web tool Onlinemeta (https://smuonco.Shinyapps.io/Onlinemeta). Onlinemeta can perform risk bias analyses that produce histograms and heatmaps, and perform meta-analysis for continuous variables, dichotomous variables, single-arm continuous variables, single-arm dichotomous variables, survival data, deft method, diagnostic tests, and subgroup analysis, as well as network meta-analysis for continuous variables and dichotomous variables. Compared with traditional software, Onlinemeta allows adjustment of many more user-parameters to customize plots, and is easy to operate, as it has a friendly interactive interface without requiring the use of coding or any software downloads. We hope that Onlinemeta can make meta-analysis more convenient to researcher community and boost meta-analysis related studies.

## Method

Onlinemeta is a web service constructed on Shiny framework (https://shiny.rstudio.com/), includes both User Interface (UI) and Server Function (server). For the UI part, the data frame display uses R package ‘DT’, and the introduction interface uses R packages ‘SlickR’, ‘svglite’ and ‘ShinyLP’. For the server part, the risk bias analysis uses R packages ‘ComplexHeatmap’, ‘robvis’, ‘colourpicker’, ‘colorspace’, ‘ggpubr’, and ‘RColorBrewer’, while meta-analysis uses the R packages ‘meta’, ‘metamisc’, ‘mada’, ‘metawho’ and ‘multinma’ for network meta-analysis. The whole interface optimization uses R packages ‘ShinyJS’, ‘ShinyDisconnect’, ‘ShinyFeedback’, ‘shinycssloaders’, ‘shinyWidgets’, ‘shinyBS’ and ‘Reactable’, ‘shinycssloaders’. For data processing, the R packages used include ‘data.table’, ‘reshape2’, ‘plyr’, ‘jsonlite’, ‘XML’ and ‘dplyr’. The images can be exported as either PDF or PNG files. Onlinemeta can be found at the address https://smuonco.Shinyapps.io/Onlinemeta/.

## Results

### Introduction

This web page introduces the main functions of new meta-analysis tool Onlinemeta, with example output graphs of risk bias analysis and meta-analysis shown in Figure 1. The version information and manager contact information are included below, and users are invited to give feedback or ask questions if necessary.

**Figure 1.**
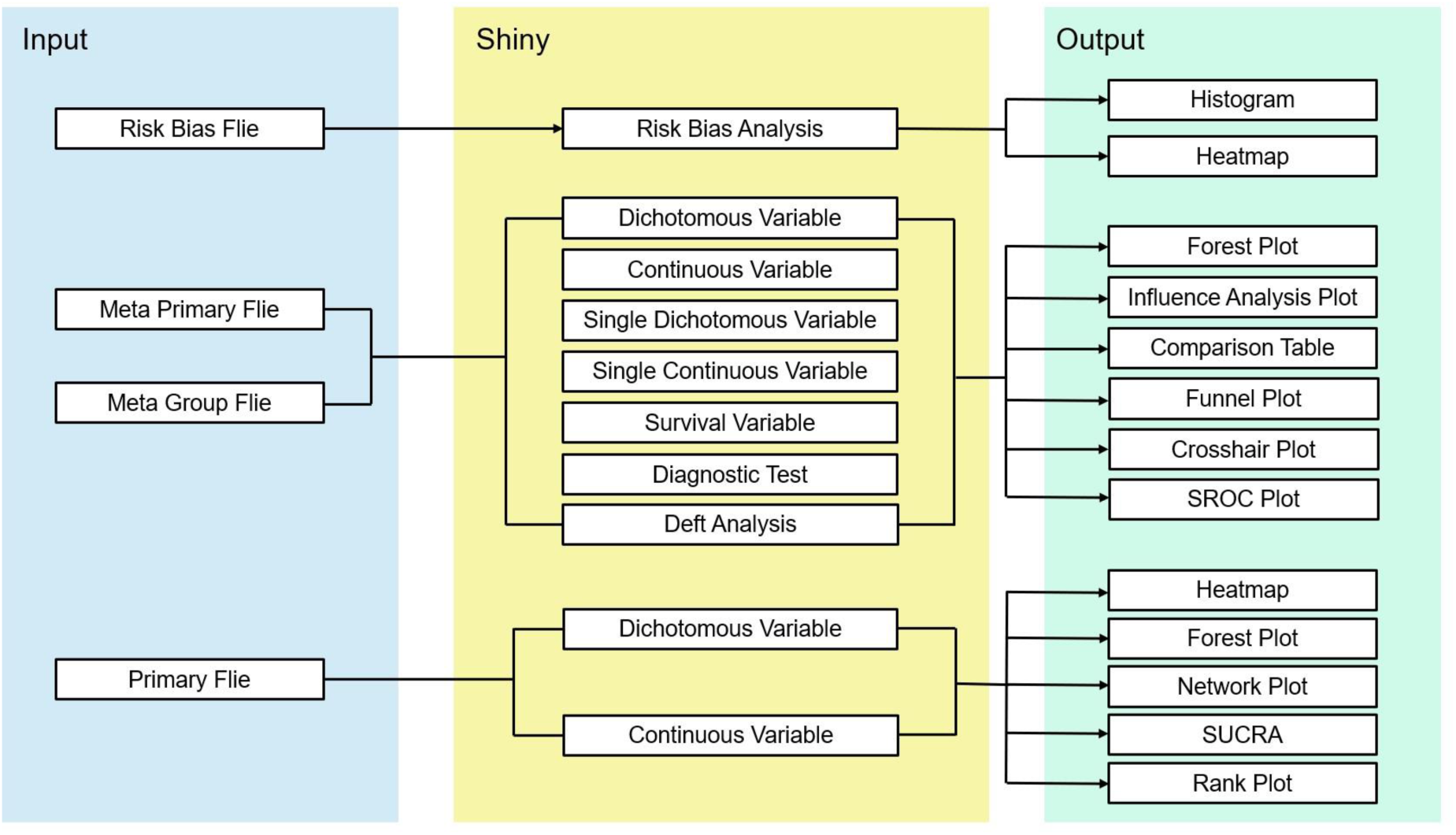
Schema describing data processing and data display for Onlinemeta.

### Histograms for risk bias analysis

This analysis feature is able to produce risk bias histograms (Figure 2). Sample data is displayed by default (Figure 2A). Users can click “Data Format” to view file formats, click the question mark behind parameters to view the explanation of the parameter, and click “Download Example Data” to download the sample data. After canceling the sample data, users can begin analysis by uploading local data to the “Upload” panel and adjusting the corresponding parameters in “Customize” panel. A series of parameters can be used to adjust plots, such as title, font size, color scheme, and size of output image (Figure 2D). The histogram shows the quality of the studies being analyzed (Figure 2B).

**Figure 2.**
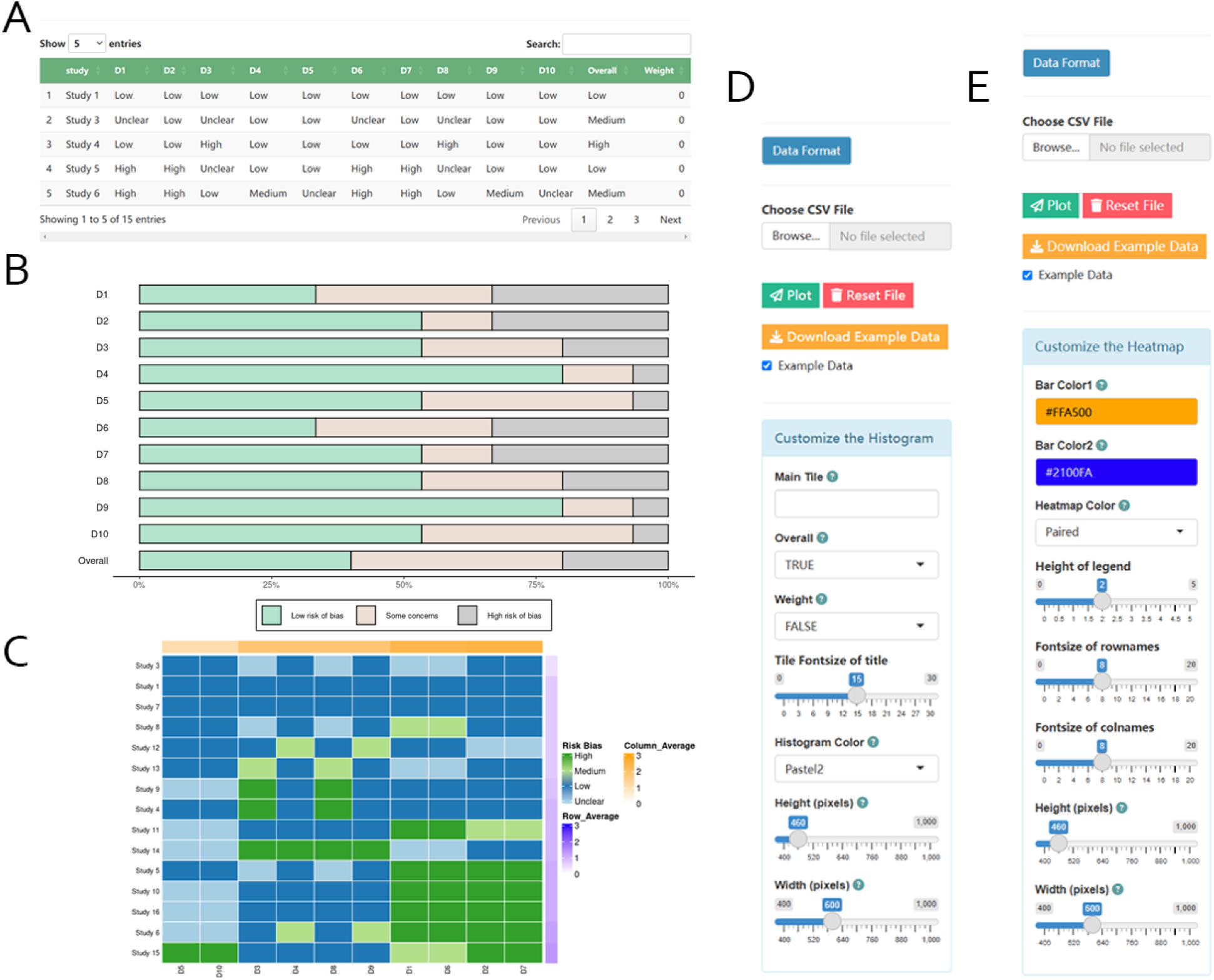
Using Onlinemeta to generate histograms and heatmaps of risk bias analysis. (A) Panels that show uploaded data. (B) A histogram generated using Onlinemeta indicating literature quality. (C) Panels for data uploading and parameter configuration to generate histograms. (D) A heatmap generated using Onlinemeta indicating literature quality. (E) Panels for data uploading and parameter configuration to generate heatmaps.

### Heatmap for risk bias analysis

Onlinemeta can output the results of risk bias analysis as a heatmap (Figure 2). Sample data is displayed by default (Figure 2A). The heatmap color represents the risk, with the labels “Unclear”, “Low”, “Medium”, and “High” assigned to 0, 1, 2, and 3, respectively. The color bars in the upper right show the average risk scores of the ranks (Figure 2C). After canceling the sample data and uploading local data in the “Upload” panel, users can adjust the heatmap color scheme, bar color, annotation height, font size, and output image size in the “Customize” panel (Figure 2E).

### The Meta function for different kinds of variables

The Meta function allows users to conduct a series of meta-analyses (Figure 3). Depending on the type of original data being analyzed, users can select “Meta” for dichotomous variables, continuous variables, single dichotomous variables, single continuous variables, and survival variables from the interface. Onlinemeta can then output four types of diagrams: forest plots (Figure 3A), funnel plots (Figure 3B), sensitivity analysis forest plots (Figure 3C), and comparison table of effects (Figure 3D). In the “Customize” panel, fixed effect or random effect models can be selected. We provide “MD”, “SMD”, and “ROM” for continuous variables, and “RR”, “OR”, “RD”, “ASD”, and “DOR” for dichotomous variables. In addition, “PLOGIT”, “PAS”, “PFT”, “PLN”, and “PRAW” are available for single dichotomous variables, whereas “MRAW” and “MLN” are available for single continuous variables. Finally, “RD”, “RR”, “OR”, “ASD”, “HR”, “MD”, “SMD”, and “ROM” can be used for survival variables. Users can adjust the label name, square color, CI color, and font size to beautify their forest plot (Figure 3E). After providing a group file, subgroup analysis can also be implemented.

**Figure 3.**
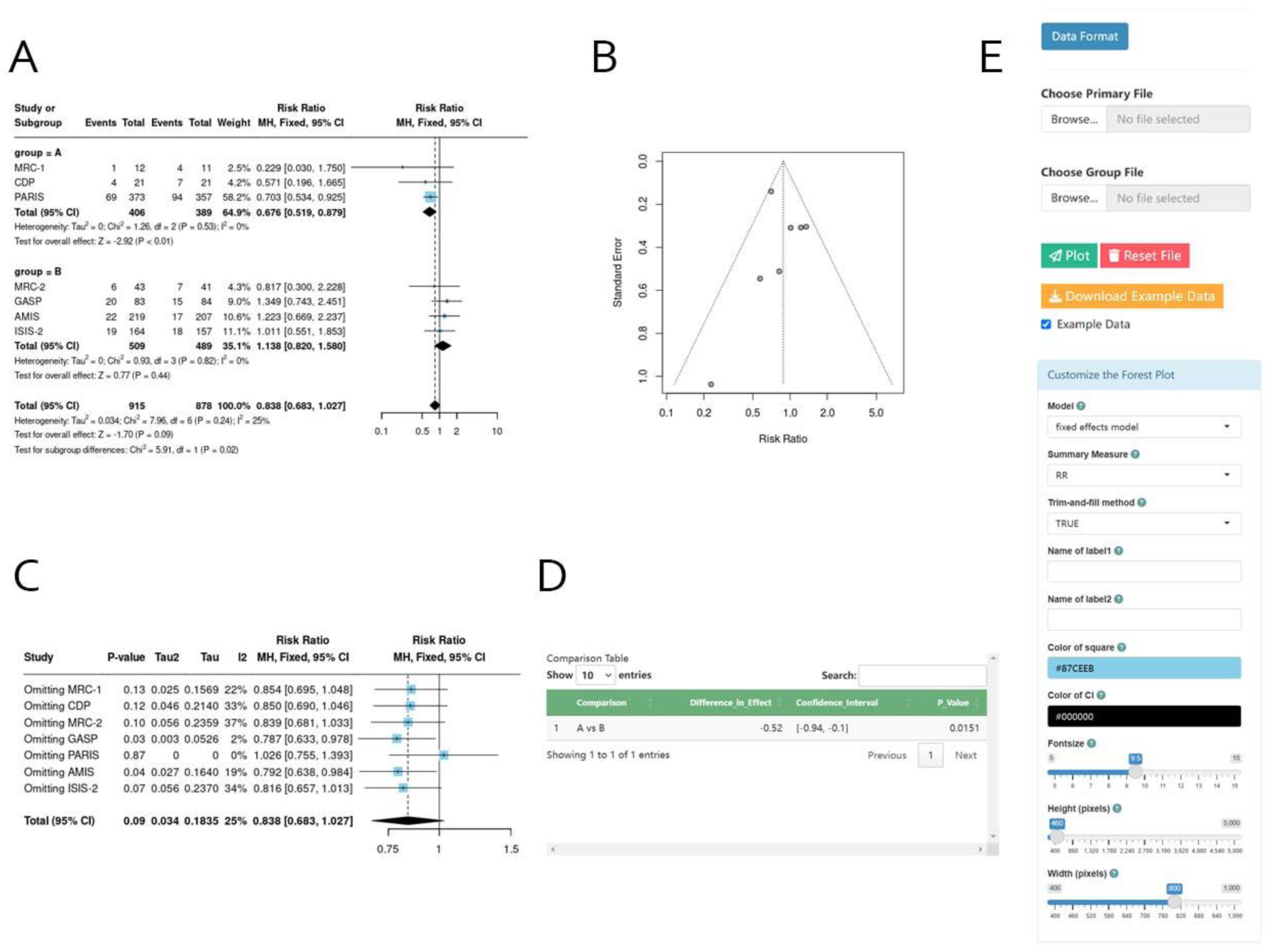
Using Onlinemeta to conduct meta-analysis for different kinds of variables. (A) A forest plot generated using the Onlinemeta showing the result of meta-analysis. (B) A funnel plot generated using the Onlinemeta showing risk bias. (C) A sensitivity analysis forest plot generated using Onlinemeta showing the the sensitivity results of the study. (D) A comparison table generated using Onlinemeta showing the comparative relationship of effects between subgroups. (E) Panels for data uploading and parameter configuration to generate forest plots.

### Onlinemeta for diagnostic tests

Users can use functionality provided by this page to conduct meta-analysis for diagnostic tests (Figure 4). Here the results have been visualized as a forest plot (Figure 4A), SROC curve (Figure 4B), and Crosshair plot (Figure 4C), demonstrating the sensitivity and specificity of both the individual research results and the comprehensive results. A series of parameters such as line color, line width, line position, line type, and symbol can be adjusted (Figure 4D).

**Figure 4.**
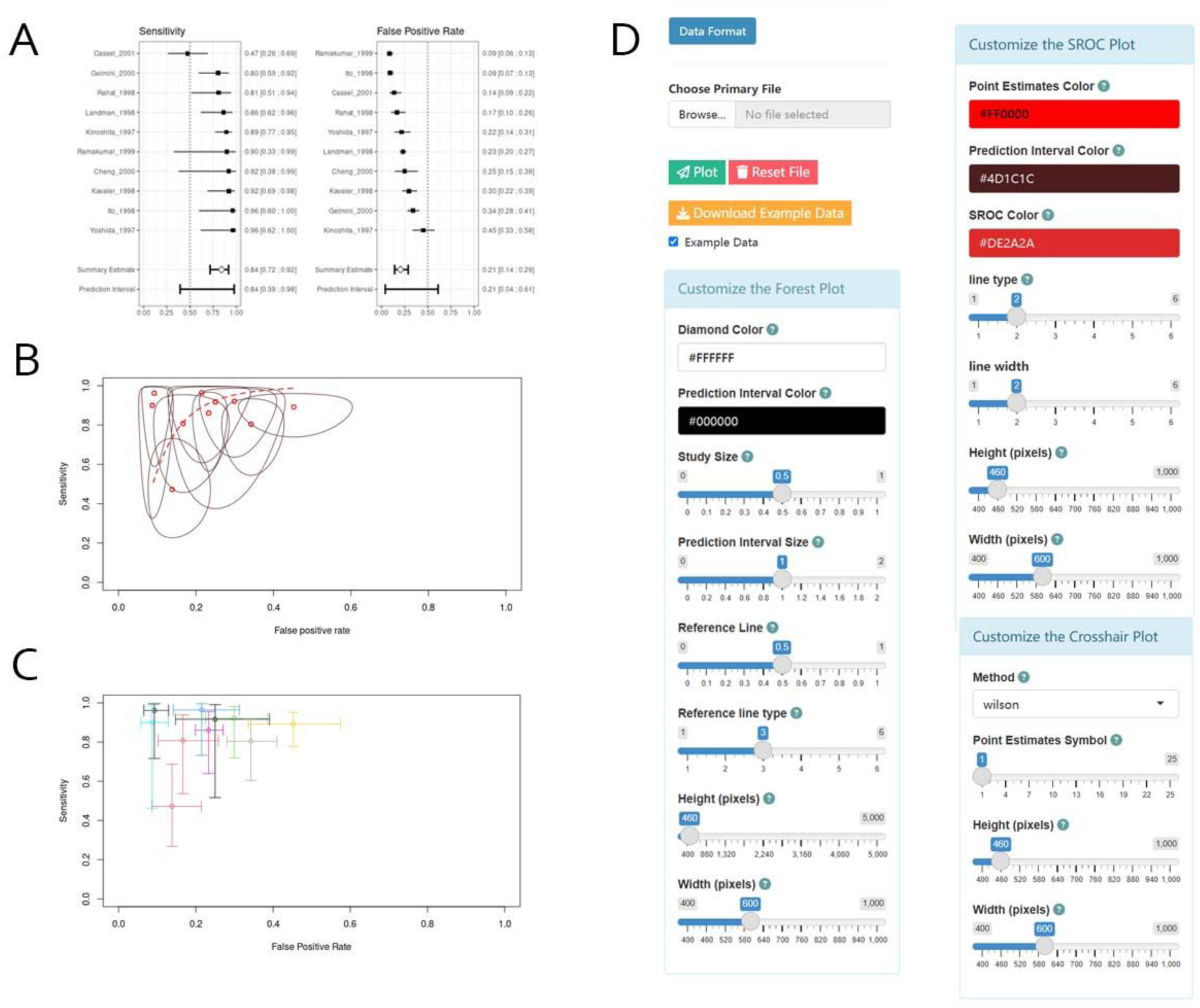
Using Onlinemeta to conduct meta-analysis for diagnostic tests. (A) A forest plot generated using the Onlinemeta showing the results of meta-analysis. (B) A SROC curve generated using the Onlinemeta showing sensitivity and specificity. (C) A crosshair plot generated using the Onlinemeta showing sensitivity and specificity. (D) Panels for data uploading and parameter configuration to generate forest plots, SROC curves, and crosshair plots.

### Onlinemeta for deft method meta-analysis

Users can use the functionality provided on this page to conduct meta-analysis using the deft method (Figure 5). The deft method follows the basic principles of meta-analysis, but compared to other methods, in meta-analyses using the deft method, the effects it focuses on are assessed based on the actual measurements and reported results in each relevant experiment, making the results more scientific and standardized[10].

**Figure 5.**
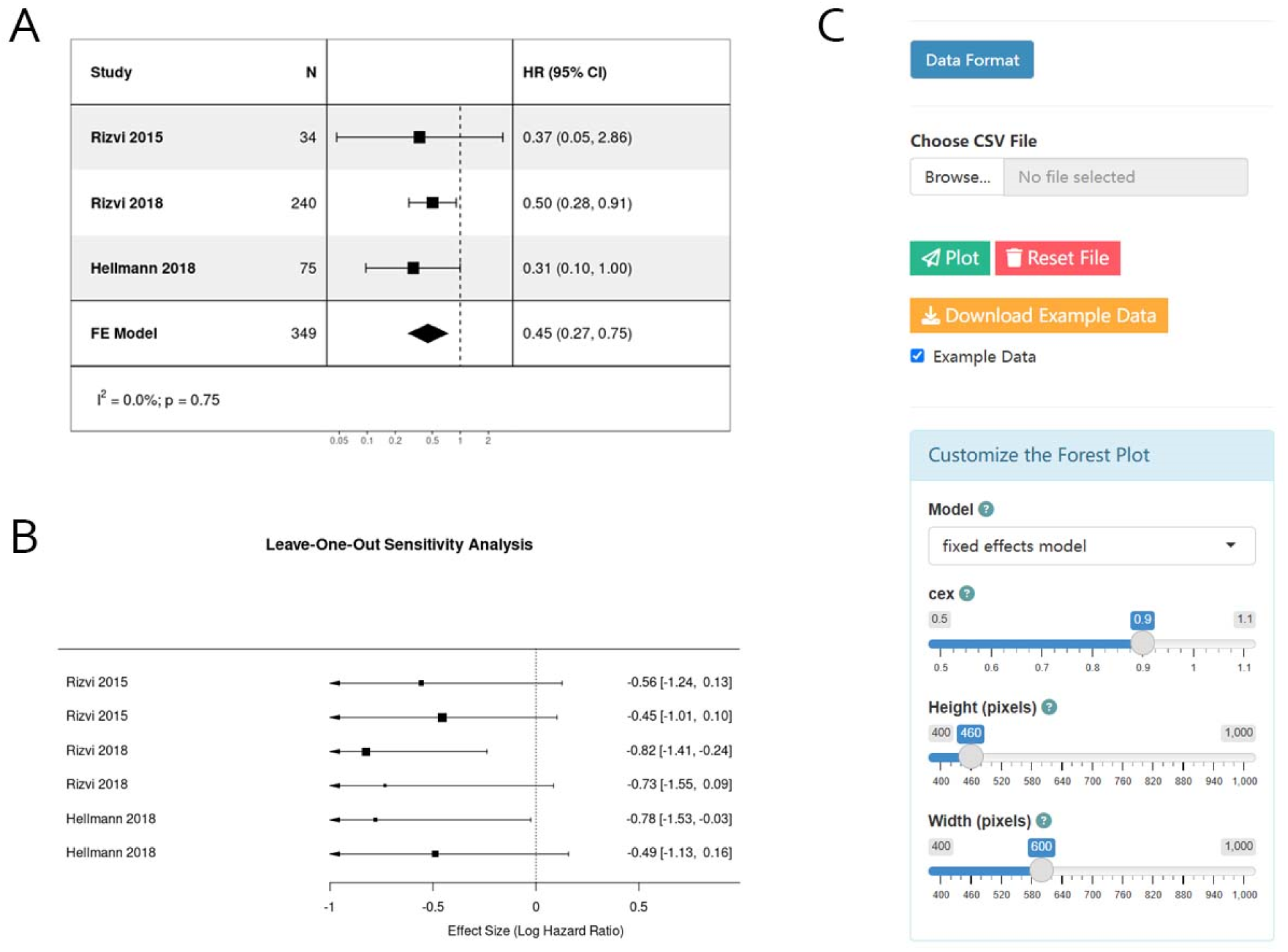
Using Onlinemeta to conduct meta-analysis in deft method. (A) A forest plot generated using the Onlinemeta showing the results of meta-analysis in deft method. (B) A sensitivity analysis forest plot generated using Onlinemeta showing the the sensitivity results of the study using deft method. (C) Panels for data uploading and parameter configuration to generate forest plots.

Here the results have been visualized as a forest plot (Figure 5A) and sensitivity analysis forest plots (Figure 5B). Users can adjust the effect model, image size, and spacing size (Figure 5C).

### Onlinemeta for network meta-analysis

Users can use the functionality provided on this page to conduct network meta-analysis (Figure 6). Here the results have been visualized as network plots (Figure 6A), forest plots (Figure 6B), SUCRA (Figure 6C), rank plots (Figure 6D), and heatmaps (Figure 6E). Directly or indirectly compare and synthesize the relative effects of multiple studies. Users can adjust a series of parameters, such as line color, line width, line position, line type and symbol (Figure 6F).

**Figure 6.**
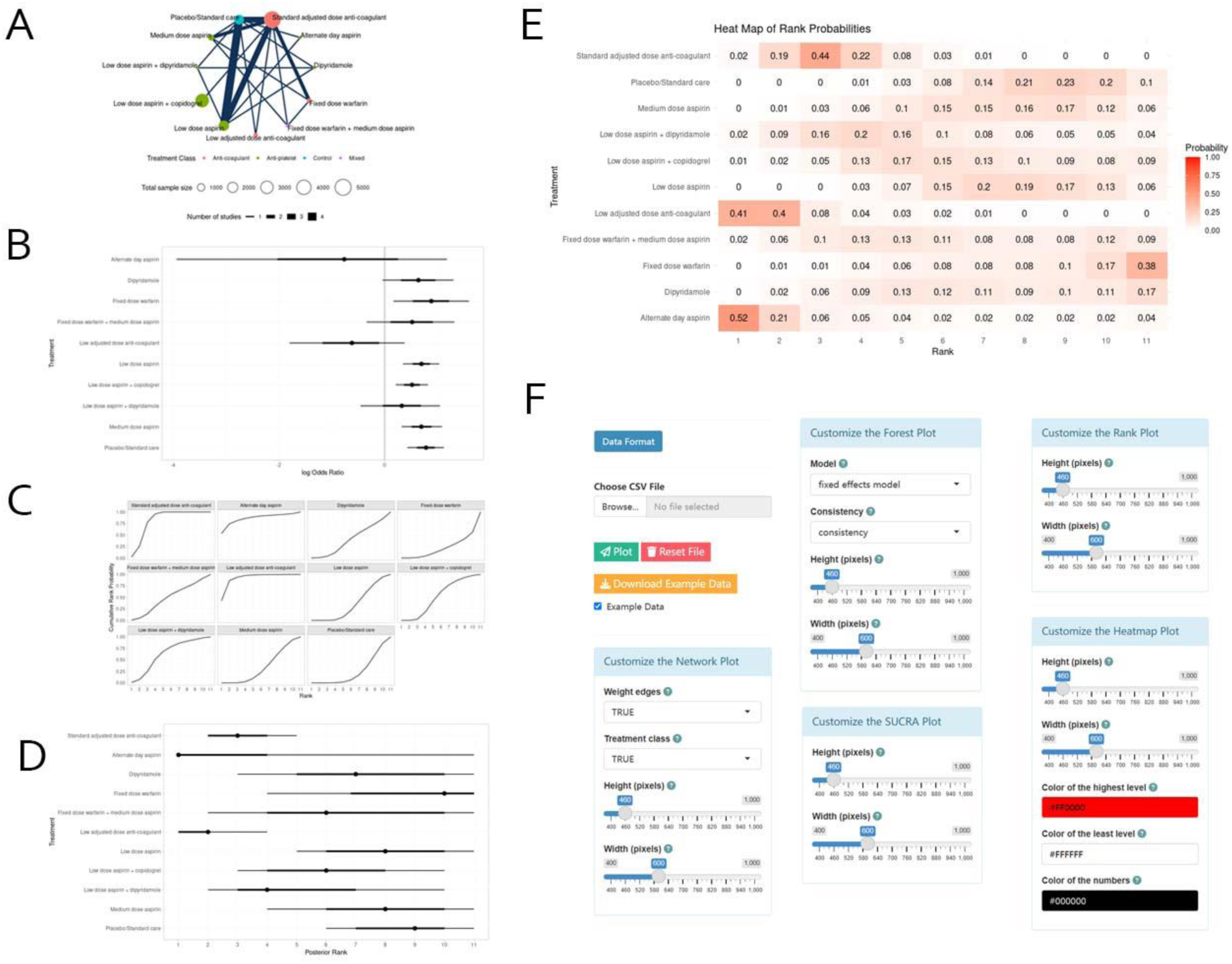
Using Onlinemeta to conduct network meta-analysis. (A) A network plot generated using the Onlinemeta showing the results of network meta-analysis. (B) A forest plot generated using the Onlinemeta showing the results of network meta-analysis. (C) A SUCRA plot generated using the Onlinemeta showing ranking probability of each study in curve plot. (D) A rank plot generated using the Onlinemeta showing ranking probability of each study. (E) A heatmaps generated using the Onlinemeta showing ranking probability of each study. (F) Panels for data uploading and parameter configuration to generate network plots, forest plots, SUCRA plots, rank plots and heatmaps.

### Help

This page provides FAQ section for solutions to common problems encountered when using this software. Additionally, it includes a detailed user manual to guide users in understanding and utilizing the tool effectively. At the bottom of the page, a dedicated email contact is provided to facilitate user feedback and inquiries.

## Discussion

Meta-analysis can combine a large number of studies from around the world, providing less biased and more accurate results than possible from single studies for potential use in fields such as evidence-based medicine[11, 12]. However, at present, most meta-analysis is performed using programming languages or software, and some of this software is often missing several useful functions [6, 7]. To fill this gap, here we presented Onlinemeta, an interactive web tool based on R-Shiny, which provides a new way to perform risk bias analysis and meta-analysis.

Onlinemeta is full featured and free accessible tool. In recent years, many online meta-analysis web tools have also been developed based on R-Shiny, such as IPD Mada[13] and MetaDTA[14]. However, our Onlinemeta is the first tool capable of performing not only risk bias analysis, seven kinds of meta-analysis, subgroup analysis and network meta-analysis simultaneously. For risk bias analysis, in addition to the traditional histogram, we designed the tool to produce a risk heatmap, which calculates the average risk score of each row and column. Users can also choose whether to include ‘overall’ and ‘weight’ as factors in the risk bias analysis. We also enabled the use of meta-analysis for dichotomous variables, continuous variables, single dichotomous variables, single continuous variables, survival variables, diagnostic tests or in deft method. In addition, users can upload group files for subgroup analysis and select the fixed effect/random effect model by adjusting the parameters.

Onlinemeta has highly customizable analytical plots. It also provides PDF and PNG files for each output image. In the risk bias analysis section, we provide a variety of color schemes for histograms and heatmaps, enabling users to choose or customize colors. The title, font size, and size of output image can also be adjusted. In the meta-analysis section, we provide users with a variety of drawing methods and models. Relevant parameters such as name, line color, line type, line width, logo, and size of output image can also be adjusted to facilitate customizing pictures. As an interactive interface, Onlinemeta is easy to use regardless of whether users have any prior statistical background or programming knowledge. In addition, directly uploading the ‘.csv’ format file avoid the need for complicated data preparation beforehand. Moreover, users do not need to download any software when using Onlinemeta, saving the user both time and storage space. However, as Onlinemeta is still in version 1.1, many functions can still be improved. We welcome any user feedback to help Onlinemeta become a more mature, friendly, and widely used meta-analysis software.

## Consent for publication

Not applicable.

## Data Availability

R is a programming language and environment for statistical computing available in https://www.r-project.org/. Shiny is an open-source collaborative initiative available in the GitHub repository (https://github.com/rstudio/shiny).

## Authors’ contributions

Writing-original draft, W.Y.T.D., Y.L.Y., A.Q.L., C.Z.Z., J.Z., S.X.W., M.S., P.L.; Conceptualization, S.X.W., M.S. and P.L.; Investigation, W.Y.T.D., Y.L.Y., A.Q.L., C.Z.Z.; Writing-review and editing, W.Y.T.D., Y.L.Y., A.Q.L., C.Z.Z., J.Z., S.X.W., M.S., P.L.; Visualization, W.Y.T.D., Y.L.Y. All authors have read and agreed to the published version of the manuscript.

## Acknowledgements

Not applicable

